# Dual Palmitoylation of PRCD, a Photoreceptor-Specific Protein Linked to RP, Alters Protein Stability and Subcellular Localization

**DOI:** 10.1101/2022.05.30.494045

**Authors:** Boyden Myers, Emily R. Sechrest, Gabrielle Hamner, Sree Motipally, Joseph Murphy, Saravanan Kolandaivelu

## Abstract

Progressive rod-cone degeneration (PRCD) is a photoreceptor outer segment (POS) disc-specific protein essential for maintaining outer segment (OS) structures, while also contributing to rhodopsin packaging densities and distribution in the disc membranes. Previously, we showed PRCD undergoing palmitoylation at the sole cysteine (Cys2), where a mutation is found linked with retinitis pigmentosa (RP) that is crucial for protein stability and trafficking to POS. PRCD has several predicted structural domains with unknown significance, such as the polybasic region (PBR) where an Arg17Cys (R17C) mutation is linked with RP. In this study, we demonstrate that a mutation in the PBR augments additional palmitoyl lipid modification observed through acyl-RAC in the mutant cysteine (R17C). Immunolocalization of transiently expressed R17C protein in hRPE1 cells depicts similar characteristics to wild type (WT); however, a double mutant lacking endogenous palmitoylation at the Cys2 position is comparable to the C2Y protein as both are likely aggregated and mislocalized in the mitochondria. Subretinal injection of C2Y, R17C, and R17C/C2Y mutants followed by electroporation in murine retina exhibit mislocalization in the inner segment compared to WT PRCD. Our results in the R17C mutant show palmitoylation transpires at two different locations. Despite being dually palmitoylated and demonstrating membrane association, the mutation in the PBR affects protein stability and trafficking to the OS. Moreover, palmitoylation within the PBR alone does not compensate for protein stability or trafficking, revealing the PBR domain is indispensable and any defects likely lead to dysregulation of PRCD protein associated with blinding diseases.

---

Retinitis pigmentosa (RP) is a group of inherited blinding diseases characterized by progressive retinal degeneration affecting rod photoreceptors initially leading to peripheral cell death, followed by cone photoreceptor death and ultimately complete loss of vision (1-3). To date, over 90 genes are associated with RP, including photoreceptor outer segment disc membrane protein “progressive rod-cone degeneration” (PRCD) (3-6). PRCD has been shown to be associated with RP in humans and dogs, causing late-onset degeneration of retinal photoreceptor cells (5). There are six mutations in PRCD linked to RP, with the most common mutation being “cysteine 2 tyrosine” (C2Y) found in over 30 dog breeds and in humans (5,7-11). The *Prcd* gene is expressed exclusively in retinal photoreceptor cells and encodes for a small 54 amino acid (aa) protein in humans and canines, and 53 aa protein in mice (Fig. 1A). PRCD is more concentrated at the base of the photoreceptor (PR) outer segment (OS) playing a critical role in OS disc morphogenesis (5,11,12). Earlier, our studies showed that PRCD lacking retina leads to defects in rhodopsin packaging density and distribution in the disc membranes (11). However, the connection between PRCD and rhodopsin packaging density in the disc membrane is not clearly understood. As PRCD interacts with rhodopsin, the importance of their interaction and the specific domains involved with the interaction is yet to be demonstrated. This information could divulge the precise role of PRCD in OS maintenance (12). As PRCD has been shown to be localized to the cytosolic surface of the outer segment (OS) disc membrane, although it does not appear to have a role in phototransduction exclusive presence in the disc membrane and its function remains widely unknown (11-13). Since the structure of the PRCD protein is not known, understanding the exclusive presence in the disc membrane is imperative. However, PRCD is predicted to have a few structural domains, including an N-terminal transmembrane helix followed by a cluster of polybasic residues (5,7). Previously, our lab has shown that PRCD is palmitoylated through a post-translational lipid modification on its sole cysteine, where the most common mutation (C2Y) is associated with loss of this palmitoyl lipid modification leading to severely reduced protein stability and mistrafficking to the photoreceptor inner segment (IS) (5,7,11,14). Interestingly, while this lipid modification is important for PRCD localization and stability, it does not affect membrane association (7,12). Based on this information, the speculated structural domains of PRCD, which enhance its association with disc membranes, are indispensable for POS maintenance and disc stacking.

**FIGURE 1.**
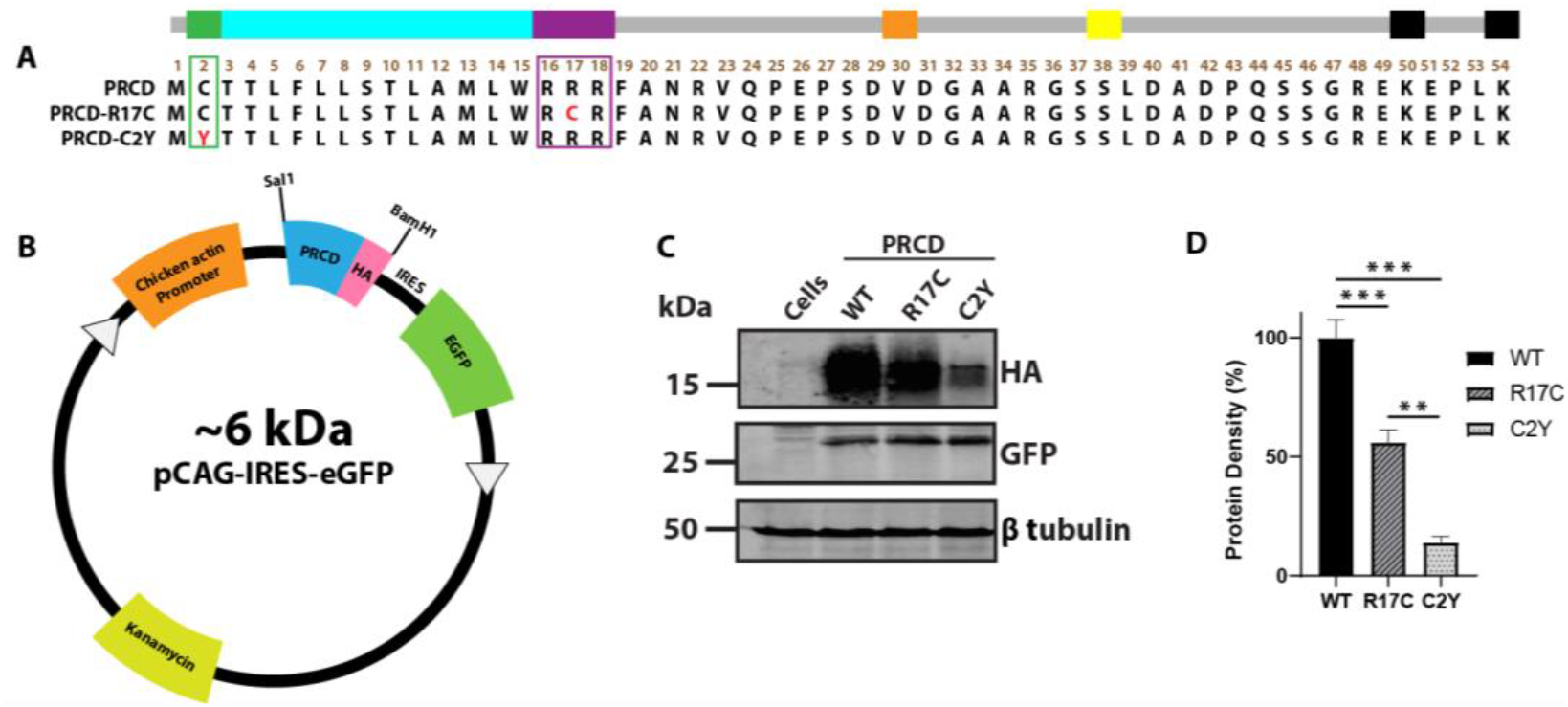
Polybasic region in PRCD is essential for Protein Stability. *A*, amino acid sequence of *PRCD* protein depicting *PRCD-WT, PRCD-R17C* and *PRCD-C2Y*. Green area indicates site of palmitoylation at the second cysteine residue, and most common mutation “Cys2Tyr” is indicated in red. Purple box depicts the polybasic region from the 16-18 aa and R17C mutation is shown in red. The transmembrane helix is coded blue from the third to fifteenth aa. Orange is the site of a valine to methionine mutation. Yellow is a phosphorylation site. The two black boxes at aa 50 and 54 are sites of ubiquitination. *B*, cloning cassette scheme demonstrating the PRCD wildtype and mutants tagged with HA (hemagglutinin) expressed under a chicken actin promoter. Also, eGFP expressed independently under IRES (Inter Ribosomal Entry Site) serves as an internal control. *C*, immunoblot of *PRCD-WT, R17C* and *C2Y* transiently transfected in hRPE1 cells. GFP serves as an internal control, with β tubulin being a loading control. *D*, quantification of *PRCD* proteins identified in *C* (n=3). The expression of *PRCD* was normalized to the internal GFP control (**, p<0.005 and ***, p<0.0005) based on a one-tailed Student t-test.

In the present study, we evaluated the importance of the polybasic region (PBR), where there are multiple mutations linked with the RP region from aa 16-22 (R17C, R18X) (5,9). Additionally, a homozygous nonsense mutation (R22X) in three siblings in a small Arab village causes RP, emphasizing the importance of the highly conserved N terminal region of PRCD (8). Many mutations in PRCD observed in humans are located in this highly conserved first 24 amino acids, coded from exon 1. Further understanding of the PBR, the mutations associated with RP, and its role in PRCD’s function and trafficking to the OS are indispensable.

Based on this information, we believe there could be molecular components in PRCD involved in the disc membrane association for maintaining OS structure. For these reasons, the molecular components could be a transmembrane helix, lipid anchor, and/or a PBR. Due to discs being double membranous structures, the need for core hydrophobic domains, lipid moiety, and polybasic residues could play a vital role in their structure and function. Therefore, in the photoreceptor cells, many of the phototransduction proteins are either attached to transmembrane domains or with specific lipid moieties (15-18). We speculate the exclusive presence of PRCD protein in the disc membranes could be a candidate for maintaining OS structure disc stacking. As identified in small GTPase Ras and Rac1 proteins, the C-terminal region contains dual lipid modifications along with poly-basic residues, which are vital for keeping a strong membrane association and efficient trafficking (19-23). Also, the G-protein transducin alpha subunit contains N-terminal lipid modifications, along with transmembrane helices, crucial for membrane attachment and for efficient function (24). Similarly, prenyl-lipid modification in the transducin gamma subunit is essential for its ciliary trafficking to the photoreceptor OS despite being associated with its partner beta subunits (25). Although PRCD does not have dual lipid modifications, perhaps the additional transmembrane helix and the PBR are key components for the strong disc membrane association. Based on this information, we hypothesized the PBR region in PRCD, along with the transmembrane helix, enhances membrane binding which is crucial for trafficking to the POS. To understand this, we tested one of the mutations associated with RP in the polybasic region (R17C). Interestingly, we find the mutant cysteine in the PBR (R17C) undergoes additional palmitoyl lipid modification, which does not affect the membrane anchor. However, despite being dually palmitoylated in the Cys2 and Cys17 positions, protein stability and efficient trafficking to the POS are significantly affected, and likely cause for RP phenotypes observed in human patients.

## Results

### Mutation in the polybasic region (R17C) significantly destabilizes PRCD protein

To determine the role of the polybasic region (PBR) in PRCD protein which is linked with retinitis pigmentosa in humans (Fig. 1*A*), we cloned gBlock gene fragments of human wild type and mutant *PRCD* constructs (*PRCD*-WT; *PRCD*-R17C; and *PRCD*-C2Y) tagged with Hemagglutinin (HA) in the C-terminal region under the control of a chicken β-actin promoter encoded by two restriction enzymes: Sal1 and BamH1 (5,7) (Fig 1*A*, and *B*). Several studies have shown the importance of the PBR in membrane association maintained through electrostatic interactions with the acidic phospholipids in the plasma membrane through the promotion of protein trafficking and protein-protein interaction. We determined to study the role of the PBR in PRCD protein (19,27-30). Previous studies, including ours, demonstrated the strong association of PRCD to the cellular membrane despite the lack of palmitoylation (7,11-13). To better assess the role of the PBR in PRCD protein stability and expression, we used GFP protein as an internal loading control for PRCD, which is driven independently by an internal ribosome entry site (IRES) (Fig. 1*B*). As we previously demonstrated, the PRCD WT and mutant plasmid constructs were transiently transfected into hRPE1 and HEK-293T (data not shown) cells to measure the stability of PRCD protein (7). Immunoblotting analysis revealed that a mutation in the polybasic region (R17C) affects protein stability by 45% (p=0.001) compared with WT PRCD expressed in hRPE1 cells. As demonstrated in our previous study, the stability of HA-tagged PRCD-C2Y protein is severely affected by >90% (Fig. 1*C* and *D*). β-tubulin and internal GFP were used as loading controls, observing no differences in comparison between WT, C2Y, and R17C transfected in hRPE1 cells (Fig.1*C*). Overall, our results show the mutation in the polybasic region (R17C) significantly affects protein stability compared with PRCD-WT despite having an additional cysteine at the 17^th^ position. Collectively, our results depict the PBR is essential for PRCD protein stability.

### The addition of a cysteine in the polybasic region does not alter protein palmitoylation and membrane association

As the mutation in the PBR (R17C) significantly affects protein stability, we assessed the palmitoylation status and membrane association of transiently transfected PRCD-R17C mutant protein in the hRPE1cell line by acyl-RAC along with PRCD-WT, as described in our earlier studies (7). Immunoblotting post-acyl-RAC revealed a robust palmitoylation status in PRCD R17C mutant protein compared to WT PRCD (Fig. 2*A*, lanes 10 and 15). As expected, in the control (-NH_2_OH), no proteins were seen in the elution fraction (Fig. 2*A*, lanes 8 and 13). GFP protein served as an internal control and was not observed in the elution fraction with +HA, suggesting PRCD-WT and R17C are both palmitoylated despite having a mutation in the PBR and severe protein instability. Ubiquitously expressed GAPDH shown in the elution fraction as palmitoylated served as a positive control (Fig. 2*A*, lanes 5, 10, and 15). We wanted to further evaluate the PBR’s role in membrane association since the PBR in Ras, and other proteins enhance strong membrane association. Additionally, as PRCD strongly binds with membranes despite having a mutation in cysteine (C2Y), we speculate the transmembrane helices and adjacent PBR are likely playing a significant role in the membrane anchor. We performed membrane fractionation and evaluated immunoblotting results to understand the PBR as a membrane anchor. Our data shows a mutation in the PBR does not affect the membrane association and the majority of proteins remain strongly associated with the membrane (>90%), similar to WT and C2Y PRCD proteins (Fig. 2*B*, lane 9; and *C*). A known membrane-binding protein, calnexin, which strongly associates with the membrane, served as a positive control (Fig. 2*B*). GFP serving as an internal control is exclusively present in the soluble cytosolic fraction (Fig. 2*B*). Overall, our data demonstrates the PRCD mutation in the PBR, linked with the RP, neither affects palmitoylation nor disrupts membrane association.

**FIGURE 2.**
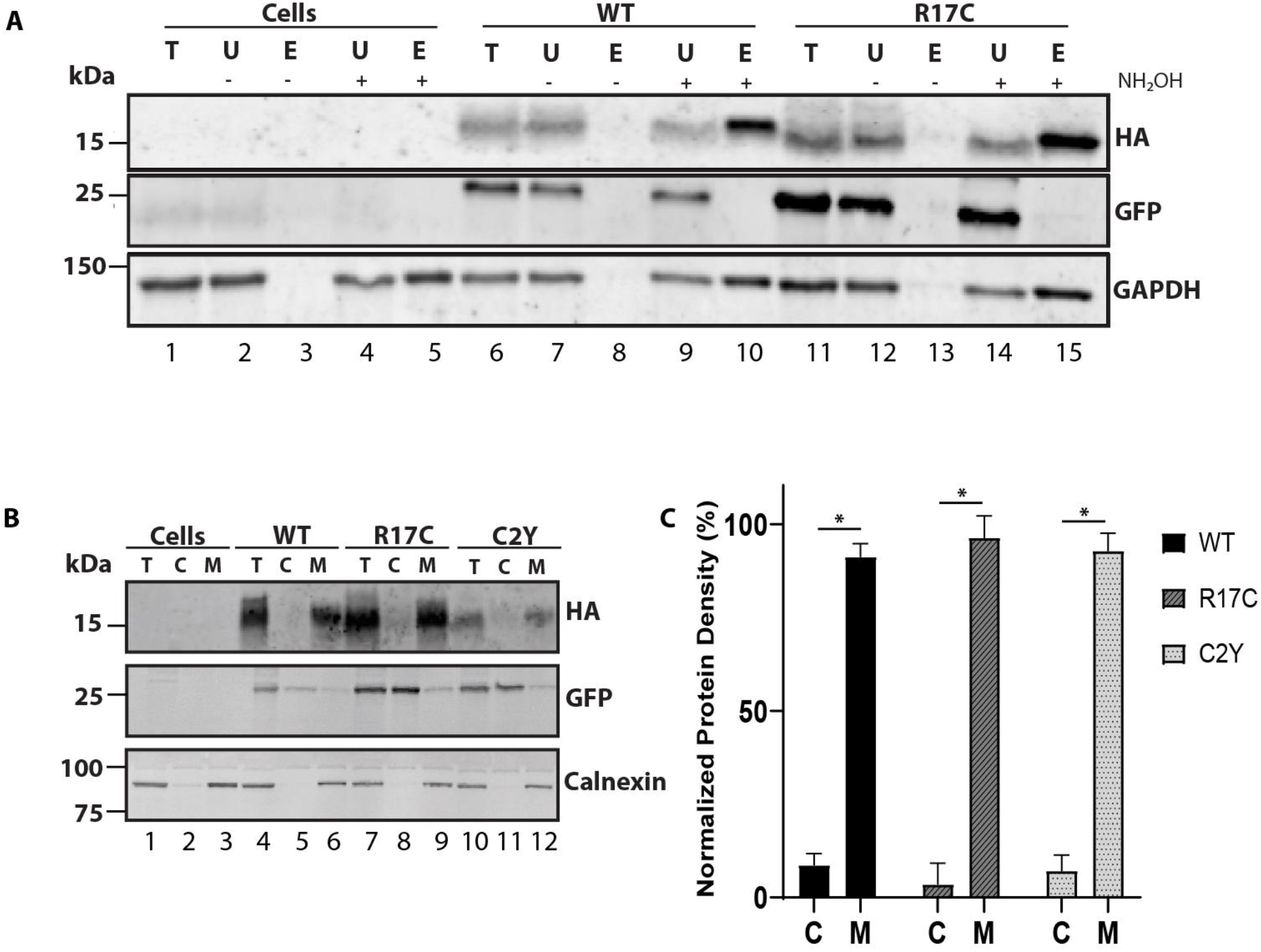
Mutation in the polybasic region of PRCD does not affect protein palmitoylation and membrane association. *A*, to assess the palmitoylation status of *PRCD-R17C*, Acyl-RAC method was conducted with protein transiently expressed in hRPE1 cells. Immunoblots show both PRCD-WT and R17C are palmitoylated after treating with hydroxylamine (+NH_2_OH) (compare lanes 8 and 10 (WT); 13 and 15 (R17C)), whereas -NH_2_OH extracts were treated with vehicle control of water (N=3). Total (T), unbound (U), elute (E) protein fractions, -E (untreated with NH_2_OH), +E (treated with NH_2_OH). GFP serves as a negative control, and GAPDH serves as a known palmitoylated positive control. Non-transfected cells (lanes 1-5) are used as a control which do not express HA tagged PRCD proteins. *B*, isotonic protein extraction examining the associations of proteins with membranes. After subcellular fractionation of hRPE1 cells transfected with indicated PRCD proteins, total (T), cytosolic (C) and membrane (M) fractions were analyzed by immunoblots with indicated antibodies. HA, PRCD protein tagged with HA-tag, GFP, a known cytosolic protein independently expressed under IRES in the same plasmid construct, and calnexin, a known membrane bound protein, were used as controls. Each PRCD mutant protein shows strong membrane binding. *C*, Protein density of each fractions were measured using Li-COR Odyssey infrared imager and normalized to protein density found in the total protein fractions (n=3) with each mutant being significantly (*, p>0.05) bound to the membrane.

### Mutant cysteine in the PBR is palmitoylated and does not rescue protein stability despite having a strong membrane association

To understand the strong membrane association of PRCD protein and the importance of the PBR, we examined the palmitoylation status of the mutant cysteine in the 17^th^ position (R17C) by mutating the sole cysteine palmitoylation site to create double mutations in PRCD (PRCD-R17C/C2Y). Our evaluation of potential palmitoylation status by palm prediction software (CSS-palm) reveals a high probability of palmitoylation in the mutant cysteine at the 17^th^ position (Fig. 3*A*) along with the known palmitoylated cysteine. The amino acid sequence of the mutant construct made in the pCAG-IRES-eGFP vector shown in Fig 3*B*. The transiently transfected PRCD double mutant shows a significant loss of protein stability compared to PRCD-WT (Fig. 3*C* and *D*). However, the stability of the PRCD double mutant protein is better than the palmitoyl lacking PRCD-C2Y protein. We showed in our earlier study that PRCD-C2Y mutation affects protein stability by 95% compared to WT PRCD. In contrast, the double mutant PRCD protein (PRCD-C2Y/R17C) is more stable (40%) compared with C2Y (5%) (Fig 1D), which suggests the cysteine in the 17^th^ position is likely palmitoylated as predicted by CSS-Palm and could stabilize the double mutant protein. Further evaluation by acyl-RAC assay demonstrates the double mutant PRCD protein is palmitoylated as predicted at the cysteine in the 17^th^ position (Fig. 3*E*, lanes 5 and 10). In contrast, the samples not treated with hydroxylamine (-NH_2_OH) observed no protein in the elution (Fig. 3E, lanes 3 and 8). This data suggests the patient mutation linked with the PBR (R17C) is likely dual palmitoylated, affecting the efficient trafficking to the photoreceptor OS. GAPDH served as a control showing the palmitoylation status (Fig. 3*E*, lanes 5 and 10). Based on these results, we believe the cysteine in the 17^th^ position is palmitoylated, which could be stabilizing the protein compared with PRCD C2Y protein.

**FIGURE 3.**
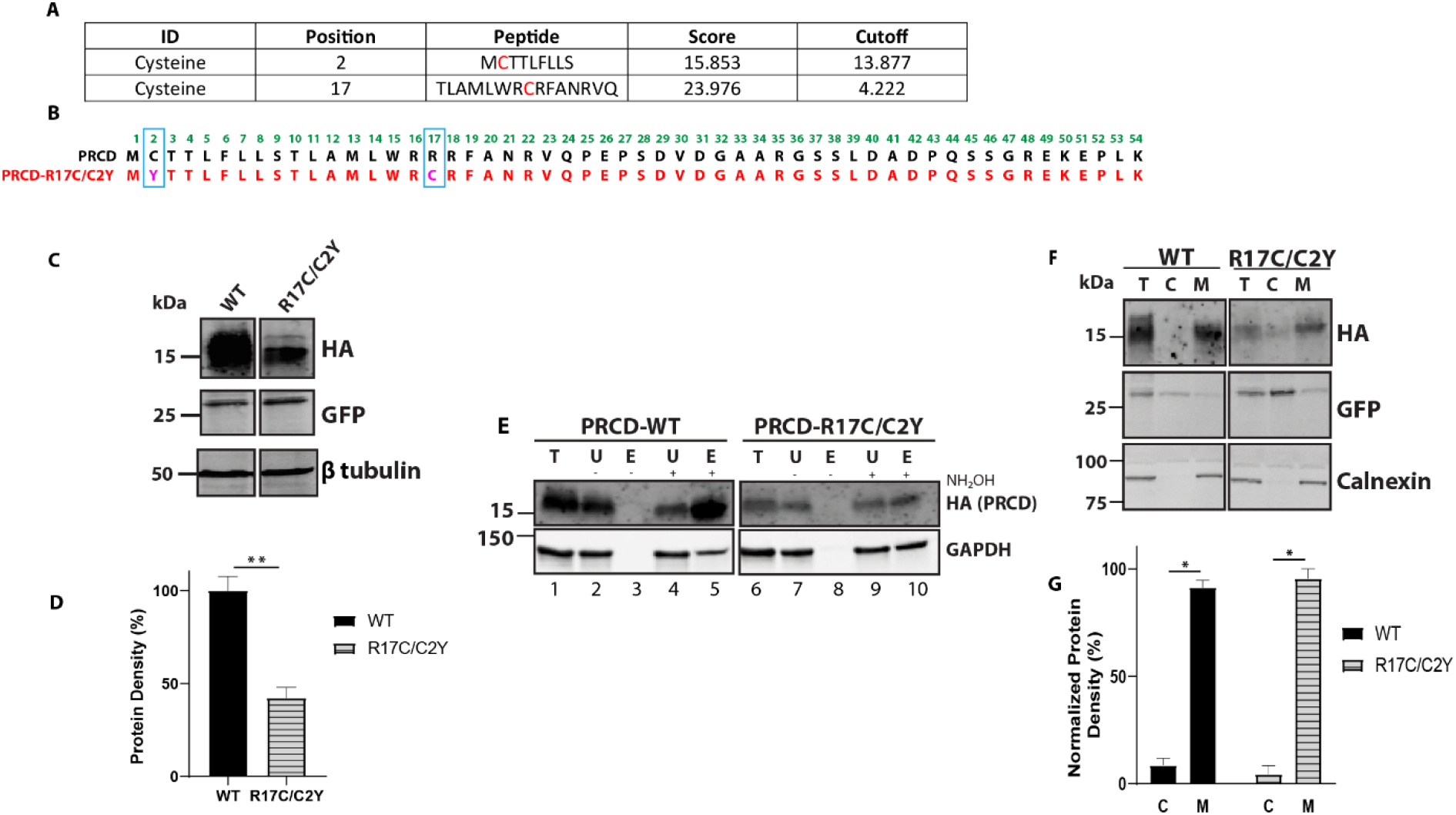
Cysteine residue of Mutant PRCD-R17C is palmitoylated and strongly binds with membranes despite the severe protein instability. *A*, CSS-Palm prediction data showing PRCD-R17C is predicted to be palmitoylated in both Cys2 and Cys17 positions. *B*, Cloning of PRCD double mutant, *PRCD-R17C/C2Y*, marked in pink with blue boxed along with PRCD-WT. *E*, Acyl-RAC of *PRCD-WT* and *PRCD-R17C/C2Y* show mutant cysteine in the 17^th^ position is palmitoylated. GAPDH serves as a positive palmitoylation control. *C*, immunoblot denoting stability of proteins: *PRCD-WT* and *PRCD-R17C/C2Y*. GFP was used as an internal loading control, with β-tubulin as an external loading control. *D*, quantification of protein stability from *panel C* (n=3, **, p>0.005). *F*, subcellular fractionation demonstrating both wildtype and mutant PRCD are found not in the cytosol (C) but is membrane (M) bound. As controls, GFP was used as a cytosolic marker and calnexin as a membrane marker. *G*, quantification of protein localization to either cytosolic fraction or membrane fraction of *PRCD-WT* and *PRCD-R17C/C2Y* (n=3, p>0.05).

In order to check palmitoylation and the PBR role in membrane association, we then performed membrane fractionation as shown in our previous study (7). Transiently transfected PRCD and mutant proteins expressed in hRPE1 cell line were performed for membrane fractionation. Immunoblotting of fractionated samples reveals a strong membrane association of PRCD double mutant, similar to WT PRCD (Fig. 3*F* and *G*). As a control, we observed no changes in solubility of GFP and membrane-bound protein calnexin fractionated in cytosolic and membrane fractions (Fig. 3*F*). Together, our results indicate that the lack of endogenous palmitoylation and the palmitoylation in the mutant cysteine is adequate for strong membrane association along with the transmembrane helices.

### Regardless of having additional lipid modification in the PBR, palmitoylation in the cysteine-2 position is indispensable for proper localization in hRPE1 cells

The exclusive localization and strong membrane association of PRCD protein in the photoreceptor OS disc membrane is essential for OS maintenance (7,11,12). Our previous study showed that palmitoylation is essential for proper localization with an unknown function because the loss of palmitoylation leads to severe protein instability (7,31). The present study demonstrates that mutation R17C localization shows similar characteristics to wild-type PRCD in hRPE1 cells (Fig.4 *A-B,E-F*). Green represents Golgi marker, GM130 (Fig. 4*A,C,E,G*), and red demonstrate mitochondrial marker, HSP60 (Fig. 4B,D,F,H). Palmitoylation deficient PRCD (C2Y) is aggregated and mislocalized in the subcellular compartments, particularly with HSP60, suggesting that palmitoylation deficiency affects efficient trafficking (Fig. 4D). However, PRCD double mutant (PRCD-R17C/C2Y), which lacks endogenous palmitoylation and mutation in the PBR, leaves R17C palmitoylation partially rescued from aggregation and localization in the subcellular compartment (Fig. 4G,H). These results suggest that despite palmitoylation in the PBR, endogenous palmitoylation is essential for proper localization. However, some mutant proteins were localized similarly to WT PRCD (Fig. 4*E,F*). As demonstrated earlier, we treated the transiently transfected PRCD in hRPE1 cells with 2-bromopalmitate (2-BP), a known palmitoyl inhibitor, for 12 hours after being transfected for 24 hours (7,32). Palmitoylation deficiency leads to mislocalization of the PRCD protein into subcellular compartments. As shown in the C2Y mutant (Fig. 4*D*), we see PRCD-WT, R17C, and R17C/C2Y proteins are aggregated and colocalized with HSP60 in the mitochondria (Fig. 4*J,N,P*). These results demonstrate loss of palmitoylation leads to PRCD protein aggregates and mislocalization to the mitochondria, which likely leads to severe destabilization, as shown in Fig. 1*C* and Fig. 3*C*. Taken together, these results unequivocally demonstrate that endogenous palmitoylation is crucial for proper trafficking since even palmitoylation in the mutated cysteine at the 17^th^ position remains mistrafficked to the mitochondria.

**Figure 4.**
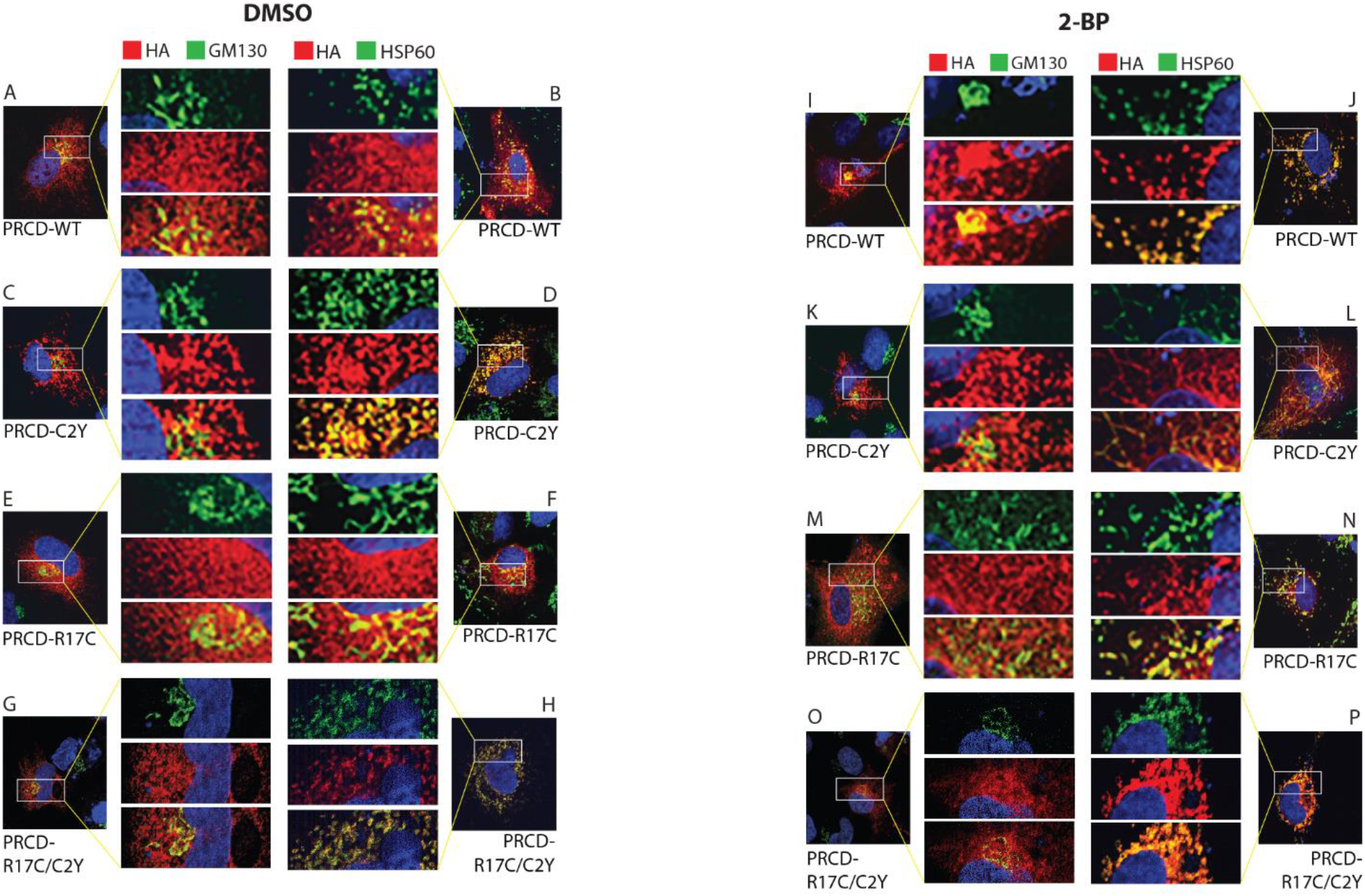
Palmitoylation in cysteine in the second amino acid position is essential for proper localization. Whereas, Immunolocalization of *R17C-PRCD* displays similar phenotype to wild type. Immunocytochemistry of transiently transfected hRPE1 cells with *PRCD-WT, PRCD-C2Y, PRCD-R17C* and *PRCD-R17C/C2Y* showing PRCD’s (HA, red) location in regards to the Golgi complex (GM130, green) and Mitochondria (HSP60, green). Nuclei are stained blue with DAPI. *A-H*, hRPE1cells treated with control, DMSO. *I-P*, hRPE1 cells treated with 150μM 2-BP for 24 hours post-transfection.

### Subretinal injection followed by electroporation in murine retina unveils PRCD-R17C mutant protein mislocalized to the inner segments

In order to reveal the localization of mutant PRCD in the PBR protein in the mouse retina, we performed subretinal injection at P0 of pups followed by *in vivo* electroporation of HA-tagged PRCD-WT, R17C, and R17C/C2Y (Fig. 5*A*). As demonstrated in our earlier study, palmitoylation deficient PRCD protein is stuck in the IS (7); so here, we examined the importance of the PBR in protein localization. Our result shows retinal cryo-sections expressing PRCD-WT in the photoreceptor OS (Fig. 5*B*) as previously reported (7),. In contrast, GFP used as a control, is expressed in the same construct driven independently under an IRES is shown in the IS as demonstrated in previous studies (Fig. 5*B*) (7,33,34). The magnified image shows the localization of PRCD-WT protein in the OS and GFP in the IS(Fig. 5*B*, enlarged).

**Figure 5.**
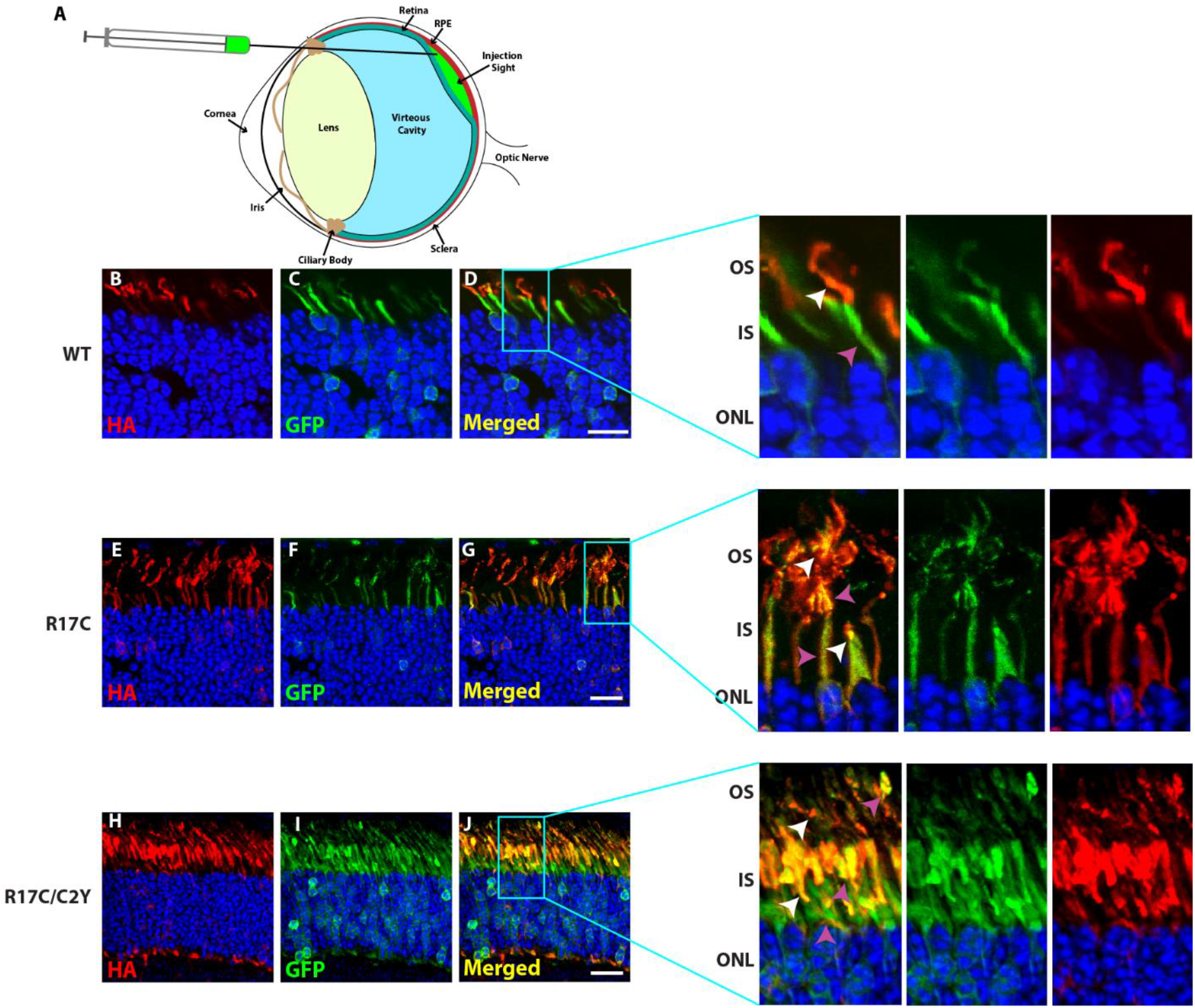
Additional palmitoyl lipid modification severely mislocalized despite being localized to photoreceptor OS. *A*, Illustration depicting site of injection in the subretinal compartment between the retina and RPE. *B-J*, Subretinal injection followed by electroporation of HA tagged PRCD-WT (*B-D*), R17C (*E-G*), and R17C/C2Y (*H-J*) double mutants in wild type mouse retina show localization of PRCD-WT in the photoreceptor OS. In contrast, PRCD-R17C is localized both in the OS and IS, and R17C/C2Y is densely mislocalized in the IS than the OS. GFP (green) serves as a transfection control localized in the photoreceptor IS as described in earlier studies. Right panels are magnified to show the localization patterns of PRCD proteins. Wild type PRCD is exclusively expressed in photoreceptor OS, and R17C show both mislocalization in the IS and normal localization in OS, whereas, majority of double mutant PRCD-R17C/C2Y protein is mislocalized to the inner segment.

Interestingly, the PBR mutant protein (R17C), which has an additional lipid modification, is localized with both the OS and IS of the photoreceptor cells (Fig. 5*G*). Though we observed the PBR mutant localization in the OS, the majority of protein is mislocalized in the IS. In contrast, the PRCD double mutant (PRCD-R17C/C2Y) protein lacking endogenous palmitoylation and containing a mutation in the PBR leading to palmitoylation of Cys17 position remains stuck in the IS (Fig. 5J). However, minor localizations are observed in the OS, likely due to the palmitoylation in the PBR region. Overall, these results demonstrate that despite palmitoylation occurring in the PBR region, exclusive endogenous palmitoylation at the Cys2 position is critical for efficient trafficking to the POS. Though the stability of the PBR protein, R17C, is significantly better than the C2Y protein (Fig. 1D), it does not traffic to the OS and remains in the IS.

## Discussion

The main goal of our study was to understand the molecular determinants responsible for the strong disc membrane association of PRCD protein, essential for maintaining photoreceptor OS structure. As we demonstrated in our previous study, palmitoylation is not a primary determinant for membrane anchoring. Therefore, we tested the importance of the polybasic region (PBR) in PRCD protein linked with RP (5), where two mutations are associated with RP (5,10). We determined to study two major goals: 1) importance of the PBR in membrane anchoring, and 2) the possibility of the mutant cysteine in the 17^th^ position (R17C) forming a disulfide linkage with endogenous cysteine at the second position; which is palmitoylated, leading to protein de-stability and mistrafficking to subcellular compartments, or additional lipidation linked with RP in humans (5). Our studies clearly demonstrate that mutation in the PBR (R17C) significantly affects protein stability (Figure 1C-D) despite being strongly membrane associated. Most interestingly, the mutant cysteine at the 17^th^ position is palmitoylated, similar to the endogenous cysteine at the second position.

Protein palmitoylation is the most common lipid modification in a protein, where a 16-carbon palmitic fatty acid is attached to a cysteine residue. Unlike other lipid modifications, palmitoylation is unique because it is reversibly modified and lacks a consensus motif requirement. We have demonstrated in an earlier study that palmitoylation deficiency in PRCD leads to severe protein destabilization and mis-trafficking to the subcellular compartments in the photoreceptor IS (7,11). In rhodopsin, palmitoylation deficiency leads to visual impairment and light-induced photoreceptor degeneration (11,15,16,35). Furthermore, in small GTPase Ras protein, there are multiple lipidations in the C-terminal region along with the polybasic region essential for efficient protein trafficking to subcellular compartments and strong membrane anchoring (36-38). We demonstrate that PRCD is predicted to have several structural domains, as described in Figure 1, with unknown significance, where many of them have mutations linked with RP in humans (5,10). In our earlier studies, we show despite lacking palmitoylation PRCD remains strongly associated with the membrane. This is likely the highly conserved N-terminal hydrophobic core transmembrane helices, followed by the PBR which potentiates the strong membrane binding (7). To understand this region further, our data from the present study reveals the PBR may not be necessary for membrane anchoring. However, our results show the mutant cysteine in the PBR is potentially palmitoylated by the Acyl-RAC approach in an *in vitro* cell culture system (Figure 2A) and show that despite being palmitoylated and strongly membrane associated (Figure 2B), the stability of the PRCD-R17C mutant protein is significantly affected (Figure 1*C*).

Although it is clear the mutant cysteine does not form a disulfide formation, additional palmitoyl lipid modification does not play a significant role in stabilizing the PRCD-R17C protein. To further understand strong membrane association, we deleted endogenous palmitoylation (C2Y), but retained the PRCD-R17C mutation in the PBR (Figure 3B). We observed despite being palmitoylated at the 17^th^ position (Figure 3*E*), the remaining PRCD-R17C/C2Y protein is severely destabilized (Figure 3*C*) without affecting strong membrane associations (Figure 3F). As we speculated in our earlier studies, the highly conserved transmembrane helices (aa 3-15) could be a primary determinant for PRCD membrane association. Based on this information, we postulate the PBR in PRCD protein likely supports strong membrane binding, along with transmembrane helices. However, additional lipid modification in the PBR likely hinders the proper exit from the Golgi compartment as PRCD undergoes palmitoylation, which may lead to hindrance in exiting from the IS. Our reasoning stems from showing immunolocalization of R17C in hRPE1 cells behaving in a similar manner as wild-type PRCD (Figure 4*A-B,E-F*). However, mutation in the endogenous cysteine (C2Y) linked with RP is aggregated and mis-trafficked to the sub-cellular compartments, particularly to the mitochondria where we observe colocalization with HSP60, a mitochondrial marker (Figure 4*C-D*). As indicated, palmitoylation of the second amino acid is likely crucial for trafficking to the OS with an unknown mechanism. Palmitoylation deficiency could also lead to defects in protein folding, which likely causes mistrafficking to the subcellular compartments where it becomes destabilized rapidly by proteolytic degradation (7). Furthermore, we speculate the mutation in cysteine could alter protein confirmation where exposed mito-signaling in the hydrophobic core alters traffic to the mitochondria.

Our *in vivo* electroporation studies show a clear discrepancy in proper localization, where the PBR mutant protein (R17C) is mis-localized to the IS (Figure 5*G*). However, with the loss of endogenous lipidation (PRCD-C2Y), palmitoylation in the PBR alone does not rescue proper localization and mutant PRCD remains ensnared in the IS (Figure 5*J*). Taken together, we suggest the PBR likely supports membrane anchoring to an extent and additional palmitoylation does not support the trafficking to the OS, likely due to improper exiting from the Golgi compartment needed for trafficking to the OS. We infer that the lack of endogenous palmitoylation and palmitoylation in the PBR could cause two potential problems that proceed to cellular dysfunction: 1) endogenous palmitoylation in Cys2 position is essential for trafficking to the POS, and 2) palmitoylation in the PBR could lead to the mutant PRCD being trapped in the IS where it is mis-trafficked and aggregated to the subcellular compartments. Based on these, despite palmitoylation in the PBR, additional cellular stress could be created due to problems in efficient trafficking, which would cause protein destabilization resulting in PRCD being trapped in the IS and eventually photoreceptor deterioration.

The overall conclusion of our study is to elucidate the effects of dual palmitoyl lipid modification (R17C) in a photoreceptor disc-specific protein, PRCD, which is linked with RP. Being the first study revealing such an effect, our findings show despite being additionally palmitoylated, proper transport from the Golgi is likely essential for proper trafficking to the POS. Any defects in this led to protein de-stability and cause additional cellular stress. Similar to the most common mutation in PRCD (C2Y), mutation in the PBR affects trafficking to the POS which could lead to the disorganization of POS disc membranes (7,11,12,14). Furthermore, our data shows that PRCD’s strong membrane association is likely due to transmembrane helices and the PBR may contribute additional support for the strong photoreceptor disc membrane association. However, the precise role of PRCD mitochondrial targeting with and without palmitoylation needs to be clarified further. As well, the cellular location of PRCD-R17C palmitoylation needs to be addressed. Endogenous palmitoylation transpires at the Golgi post-packaging through the endoplasmic reticulum (ER), but palmitoylation of PRCD-R17C/C2Y would ensue at the ER because we demonstrate it is not targeted to the Golgi. Ultimately revealing the PBR mutation, R17C, is not only dually palmitoylated but said palmitoylation occurs at two different locations.

## Experimental Procedures

### Reagents and Antibodies

For Acyl-RAC purification the following chemicals were used: Lysis Buffer (25 mM HEPES, pH 7.5 [Gibco], 25 mM NaCl [Invitrogen], 1 mM EDTA [Sigma-Aldrich], protease and phosphatase inhibitors), Blocking Buffer (100 mM HEPES, pH 7.5 [Gibco], 1 mM EDTA [Sigma-Aldrich], 2.5% SDS [Sigma-Aldrich], 0.1% *S*-methyl methanethiosulfonate [Sigma-Aldrich]), Binding Buffer (100 mM HEPES, pH 7.5 [Gibco], 1 mM EDTA [Sigma-Aldrich], 1% SDS[Sigma-Aldrich]), 2 M Hydroxylamine, pH 7.5 [Thermo Scientific], 2 M NaCl [Invitrogen] and Agarose S3 High Capacity Acyl-8;’ Capture Resin [Nanocs]. When cloning and cell culture experiments progressed, the chemicals used were: DMEM/F12 medium (10% Fetal Bovine Serum [Cytiva], 1% penicillin streptomycin [Fisher]), Trans-LT1 Transfection Reagent [Mirus], Dulbecco’s phosphate-buffered saline (1x DPBS [Corning] and 0.25% Trypsin-EDTA [Gibco], dimethyl sulfoxide (DMSO) [Sigma-Aldrich], 2-bromopalmitate [MANUFAC]. For immunohistochemistry and immunocytochemistry, the reagents used were: 4% Paraformaldehyde (16% PFA, [Electron Microscopy Sciences]) and 5% Goat Serum (Normal Goat Serum [Millipore], 0.5% Triton-X-100 [Sigma-Aldrich], 0.05% Sodium azide [MP]). Measurement of protein levels transpired with 1x phosphate buffered saline, Pierce™ Protease Inhibitor Mini Tablets, EDTA-free [Fisher], 1x phosphate buffered saline Tween-20 [Fisher], Intercept™ Antibody Diluent T20 PBS [Li-Cor], Intercept® Blocking Buffer PBS [Li-Cor]. Subretinal injection requirements included 0.1% fluorescein in PBS [AK-FLUOR].

### Animals

All handling, care, and experimental procedures involving animals were performed in agreement with National Institutes of Health guidelines. The Institutional Animal Care and Use Committee (IACUC) at West Virginia University approved the protocol used.

### Cloning and Cell Transfection

hTERT-immortalized pigment epithelial (hRPE1) cells were used and transfected with constructs encoding human PRCD, either wild type (*PRCD-WT*), *PRCD-C2Y, PRCD-R17C* or *PRCD-R17C/C2Y* mutants, with C-terminal HA epitope-tagged gBlocks gene fragments (200ng). These gene fragments were cloned into a pCAG-IRES-eGFP vector, as described in our earlier study, where PRCD and HA are under a chicken actin promoter (7). Endogenous expression of eGFP, used as an internal control, is driven independently with IRES element. The hRPE1 cells were maintained in DMEM/F12 medium enriched with 10% FBS and 1% penicillin streptomycin in a sterile incubator at 37°C with 5% CO_2_. The cells were cultured on 100 mm circular dishes and split when 70-90% confluent onto 6-well plates. For the transfection of hRPE1 Bio Mirus’s Trans-LT1 was used and the manufacturer’s protocol was followed. 48 hours post transfection, cells were collected and stored for future analysis as described below.

### Immunoblotting

Lysates of hRPE1 cells transiently transfected with PRCD protein expressing constructs, were collected as previously explained, and homogenized with 1xPBS containing protease and phosphatase inhibitors (Pierce) (7).Protein concentration was analyzed by standard BCA method with a NanoDrop spectrophotometer (ND-1000, Thermo Scientific). A 15% SDS-PAGE was used to run the samples at equal concentrations and transferred to a Immobilon-FL membrane. After transfer, the membrane was blocked with Intercept® Blocking Buffer PBS (Licor) for one hour followed by primary antibody incubation for two hours at room temperature. Antibodies from Table 1 were diluted in a 1:1 mixture of 1xPBST (Tween-20) with Intercept™ Antibody Diluent T20 PBS. Following incubation were three 1xPBST washes for five minutes each and then a 30-minutes incubation with secondary antibodies diluted in 1xPBST per concentrations mentioned in Table 2. After three washes for five minutes each with 1xPBST, the blots were scanned and densities were measured with an Odyssey Infrared Imaging System (LI-COR Biosciences) per the manufacturer’s guidelines.

**Table 1.** Primary Antibody List.

| Antibody | Dilution & Application |  | Animal Raised In | Source |
| --- | --- | --- | --- | --- |
|  | WB | ICC/IHC |  |  |
| HA tag | 1:2,000 | 1:1,000 | Rat | Sigma-Aldrich |
| GFP tag | 1:2,000 | 1:1,000 | Mouse | Protein Tech |
| β tubulin | 1:3,000 | N/A | Mouse | Sigma-Aldrich |
| GAPDH | 1:10,000 | N/A | Mouse | Protein Tech |
| Calnexin | 1:200 | N/A | Mouse | Santa Cruz Biotechnology |
| GM130 | N/A | 1:200 | Mouse | BD Sciences |
| HSP60 | N/A | 1:500 | Mouse | Protein Tech |

**Table 2.** Secondary Antibody List.

| Antibody | Dilution & Application |  | Source |
| --- | --- | --- | --- |
|  | WB | ICC/IHC |  |
| Rat 568 | 1:50,000 | 1:500 | Invitrogen |
| Rat 680 | 1:50,000 | 1:500 | Invitrogen |
| Mouse 680 | 1:50,000 | 1:500 | Invitrogen |
| Mouse 800 | 1:50,000 | 1:500 | Li-Cor |
| DAPI | N/A | 1:1,000 |  |

### Acyl-RAC Purification (Palmitoylation Assay)

The palmitoyl modification of PRCD was assessed using Acyl-RAC, as previously described (7,26). Transiently transfected hRPE1 cells were collected as described in the following. Each sample’s pellet was resuspended in 300 µl of ice-cold lysis and sonicated at 12 psi for three rounds of three seconds, being returned to ice for 30 seconds between each round. Proteins were solubilized with 1% Triton-X-100 by incubating on a nutator at 4°C for 20 minutes. Post-incubation, samples were centrifuged at 200 x g for three minutes and the supernatant of each sample was split equally into 3 tubes, 100µl in each. To block the free cysteines, 300µl of blocking buffer containing Methyl-methane Thio-sulfonate (MMTS) was added to each 100ul aliquot (3 aliquots per sample) and incubated at 40°C for 16 minutes, while being vortexed every 2 minutes. Ice-cold acetone was added to precipitate proteins during a 20-minute incubation at -20°C. Samples were centrifuged at 20,000 x g for 15 minutes at 4°C, then the supernatant was removed and the pellets were washed with 70% acetone four times. Post wash, the pellets were air dried for 30 minutes at room temperature, resuspended with 200 µl of binding buffer and sonicated to be solubilized. Each sample’s three aliquots were pooled together (600 µl total per sample) in one tube. Assay was performed for each sample treating with and without hydroxylamine (“+HAM” and “-HAM”) with 75ul of Agarose S3 High Capacity Acyl-RAC Capture Resin (Nancos) beads, prewashed with binding buffer. 45ul of sterile water was added in the “-HA” tube and 45ul of 2M Hydroxylamine (NH_2_OH, HA), pH 7.5 was added in the “+HA” tube, respectively. 250ul of each pooled sample was added to the “-HA” and “+HA” tubes and all samples were incubated on a nutator at room temperature for two hours. After the incubation, the beads were then washed four times with binding buffer. The bound fractions were eluted with 1X Lamellae sample buffer containing 10 mM DTT by boiling for five minutes and then loaded into an SDS Page Gel.

### Subcellular protein fractionation

Two protocols were used for cellular fractionation to complement and verify the result. Subcellular Protein Fractionation Kit for Cultured Cells (Fisher Scientific) was used and the protocol was executed through Step 5 to obtain the “membrane extract”. To verify, we used the same amount of collected cells per sample (3 wells of a 6-well plate) and started by suspending them in 300 µl of isotonic buffer (1 x PBS) containing phosphatase and protease inhibitors (Pierce). The cells were sonicated three times for three seconds each at 12 amps. Between each sonication, the samples were kept on ice for 30 seconds. The samples were then centrifuged at 500 x g for three minutes at 4°C. After collecting total fraction, the remaining supernatant of each sample was ultra-centrifuged at 45,000 x g for 15 minutes at 4°C. 75 µl of the supernatant was collected and each labeled as the appropriate “Soluble Fraction”. The remaining supernatant was aspirated. Each pellet was resuspended in 75 µl of isotonic buffer with inhibitors and labeled as the “Membrane Fraction’’. 5x sample buffer was added to each final fractions Total, Soluble, and Membrane tubes and boiled for five minutes before loading into an SDS-PAGE gel followed by immunoblotting.

### Immunocytochemistry

hRPE1 cells were cultured on sterile circular glass coverslips and transfected with *PRCD* constructs as previously described (7).After a 48-hour transfection, the media was aspirated and each well was washed with 1 x PBS for one minute, followed by fixation in 4% paraformaldehyde (PFA) for ten minutes and then rinsed for 30 seconds with 1 x PBS. Permeabilization was conducted with cold methanol for five minutes before three two-minute washes with ice-cold 1 x PBS. Cells were blocked with 5% goat serum for one hour at room temperature. Briefly, slides were washed with 1 x PBS before the addition of primary antibodies overnight at 4°C. The following day, each well was washed with 1 x PBS Triton-x-100 (PBST) three times for five-minutes a piece. Secondary antibody was added for one hour at room temperature. Followed by three five-minute washes with 1x PBS. Each slide was then mounted with ProLong™ Gold antifade reagent.

### Immunohistochemistry

After eyecups were sectioned onto a Superfrost Plus Microscope Slide (Fisher) a hydrophobic barrier was drawn with a PAP pen around the desired sections. The sections were washed with 1 x PBS three times for five minutes each and then were incubated with blocking buffer (1 x PBS, 10% normal goat serum, 0.5% Triton-X, 0.05% sodium azide) for one hour at room temperature. The blocking buffer was aspirated and the sections were incubated overnight at 4°C with the primary antibodies (Table 1) of choice, diluted appropriately in antibody dilution buffer (ADB) (1 X PBS, 5% normal goat serum, 0.5% Triton-X, 0.05% sodium azide). The next morning the primary antibody was aspirated and the sections were washed with 1xPBST (0.1% Triton-X-100) for 15 minutes once and twice with 1 x PBS for 15 minutes. The corresponding secondary antibody (Table 2) was diluted in ICC ADB and the antibodies were incubated on the sections for one hour at room temperature. The sections were then washed twice for 15 minutes with 1 x PBST and then repeated once with a 1 x PBS wash. After the final wash, the slides were aspirated and set out to dry for a minute. ProLong™ Gold antifade reagent (Invitrogen) was applied to the slides prior to being mounted with coverslips and left to dry for at least 24 hours before imaging.

### Subretinal injection

Plasmid DNA of *PRCD-WT, PRCD-R17C* and *PRCD-R17C/C2Y* was purified and diluted to a concentration of 5 μg/µl along with 0.1% fluorescein in 1xPBS (100 mg ml−1 AK-FLUOR, Alcon, Fort Worth, TX). On postnatal day zero to one, CD1 (Charles River) pups were anesthetized by hypothermic conditioning obtained through keeping on ice for seven to ten minutes. Using a dissecting microscope, the future eyelid was slit open to give access to the developing eye. A 30G ½ needle was used to make a hole alongside the pupil so a blunt end needle (33 gauge, 6 pk, 10 mm, 45° angle [Hamilton - Cat #7803-05]) with 0.5 µL of the DNA/fluorescein solution could be injected into the subretinal area located behind the lens. Following injection, electroporation was conducted with tweezer-like paddles (BTX model 520, 7-mm diameter) for five pulses at 80 v for 50 ms durations with 950 ms rests in between. Eyes were collected on postnatal day 21, followed by retinal sectioning and immunohistochemistry as described earlier (7).

## Statistical analysis

Data’s are expressed as means ± S.E., unless otherwise indicated. The differences between wildtype control (PRCD-WT) and Mutants PRCD (*PRCD-C2Y, PRCD-R17C, PRCD-R17C/C2Y*) were analyzed with a two-tailed student *t* test (online version, http://www.usablestats.com/calcs/2samplet).

## Acknowledgements

The members of the Kolandaivelu lab provided valuable discussion and support for this project. We thank Drs. Ramamurthy, Maxim Sokolov, and Mr. David Sokolov for their valuable suggestions and support.

## Author Contributions

S. K. conceptualization; B.M., E.S., G.H., S.M., and J.M. investigation, data analysis, development, and reviewing the manuscript. S.K., and B.M. data generation, writing, reviewing, and editing.

## Conflict of interest

The authors declare they have no conflicts of interest with the contents of this article.

### Abbreviations

The abbreviations used are:

PRCD: progressive rod-cone degeneration
OS: outer segment
IS: inner segment
POS: photoreceptor outer segment
PBR: polybasic region
RP: retinitis pigmentosa
DPBS: dulbecco’s phosphate-buffered saline
Acyl-RAC: acyl resin-assisted capture
NH_2_OH: hydroxylamine
PBS: phosphate buffered saline
OCT: optimal cutting temperature
IHC: immunohistochemistry
ICC: immunocytochemistry
WT: wild type
IRES: internal ribosome entry site
GFP: green fluorescent protein
hTERT RPE-1: hTERT-immortalized retinal pigment epithelial cells-1
2-BP: 2-bromopalmitate.

